# Genomic analyses of two Italian oyster mushroom *Pleurotus pulmonarius* strains

**DOI:** 10.1101/2020.09.03.281089

**Authors:** Guillermo Vidal-Diez de Ulzurrun, Yi-Yun Lee, Jason E. Stajich, Erich M. Schwarz, Yen-Ping Hsueh

## Abstract

*Pleurotus* mushrooms are among the most cultivated fungi in the world and are highly valuable for food, medicine and biotechnology industries. Furthermore, *Pleurotus* species are carnivorous fungi; they can rapidly paralyze and kill nematodes when nutrient-deprived. The predator-prey interactions between *Pleurotus* and nematodes are still widely unexplored. Moreover, the molecular mechanisms and the genes involved in the carnivorous behavior of *Pleurotus* mushrooms remain a mystery. We are attempting to understand the interactions between *Pleurotus* mushrooms and their nematode prey through genetic and genomic analyses. Two single spores (ss2 and ss5) isolated from a fruiting body of *Pleurotus pulmonarius* exhibited significant differences in growth and toxicity against nematodes. Thus, using PacBio long reads, we assembled and annotated two high-quality genomes for these two isolates of *Pleurotus pulmonarius*. Each of these assemblies contains 23 scaffolds, including 6 (ss2) and 8 (ss5) telomere-to-telomere scaffolds, and they are among the most complete assembled genomes of the *Pleurotus* species. Comparative analyses identified the genomic differences between the two *P. pulmonarius* strains. In sum, this work provides a genomic resource that will be invaluable for better understanding the Italian oyster mushroom *P. pulmonarius*.

## Introduction

*Pleurotus* mushrooms are among the most cultivated and consumed edible fungi in the world (Gregori *et al*. 2007; Yin *et al*. 2014). While most fungi are difficult to cultivate, *Pleurotus* can be grown commercially to high yield (Banik and Nandi 2004). In addition to their nutritional value, these mushrooms are a natural source of prebiotics (Aida *et al*. 2009) and antioxidants (Khatun *et al*. 2015), and are thus of great interest to the food industry. Also, *Pleurotus* species contain remarkable medicinal properties (Golak-Siwulska *et al*. 2018). For example, *P. pulmonarius* exhibits anti-inflammatory (Smiderle *et al*. 2008; Nguyen *et al*. 2016), analgesic and antitumor activity (Zhang *et al*. 2007). This fungus has also shown its potential in bioremediation since it is able to degrade contaminants such as aromatic pollutants (Rodríguez *et al*. 2004) and biocides (Law *et al*. 2003). More recent studies have focused on the potential outcomes for biotechnology of *P. pulmonarius*. For instance, this fungus can produce useful enzymes even while degrading waste products (Inácio *et al*. 2015) as well as being consumed after being part of biofuel production (Chen *et al*. 2020), therefore underlying the versatility of this fungus.

One of the most unknown characteristics of *Pleurotus* mushrooms is their ability to kill and feed on living nematodes (Thorn and Barron 1984) and therefore their potential for the biocontrol of parasitic nematodes (Degenkolb and Vilcinskas 2016; Sivanandhan *et al*. 2017; Castañeda-Ramírez *et al*. 2020). Although a few compounds produced by *Pleurotus* have been reported to cause paralysis in nematode worms (Kwok *et al*. 1992; Satou *et al*. 2008), a more recent study has suggested that the true identity of the toxins targeting nematodes remain to be discovered (Lee *et al*. 2020).

The advance of sequencing techniques has led to the publication of the genome of several *Pleurotus* species; however, only a few high-quality assemblies have been produced for *Pleurotus* so far. Only the latest assemblies of *P. ostreatus* (Riley *et al*. 2014; Qu *et al*. 2016; Wang *et al*. 2018) show the overall quality and contiguity necessary for more comprehensive genetic analyses of these mushrooms. In contrast most of the *Pleurotus* genomes are very fragmented, including *P. eryngii, P. tuoliensis* (Zhang *et al*. 2018), *P. platypus* and *P. citrinopileatus* (Li *et al*. 2018) ranging from 106 to 10,689 scaffolds. Additionally, other *Pleurotus* species, such as *P. pulmonarius*, have not yet been sequenced despite their importance for different industries.

We thus studied the genomes of two haploid monokaryotic strains derived from spores of a wild isolate of *P. pulmonarius* collected in Taiwan. These two haploid monokaryons (ss2 and ss5) exhibit significant phenotypic differences in their growth and toxicity against nematodes. Using PacBio long reads, we produced two high-quality annotated genomes for *P. pulmonarius*. Moreover, our assemblies represent the first available *Pleurotus pulmonarius* genomes and are among the most complete assembled *Pleurotus*. Through this report we also provide genomic comparisons between the two assemblies, identifying highly dissimilar regions that might contribute to their observed phenotypic differences. We believe that the two high quality genomes will be useful genomic resources that will contribute to a better understanding of *P. pulmonarius*, and pave the way for future comparative genomic analyses of *Pleurotus* mushrooms.

## Materials and methods

### Fungal strains and DNA extraction

Spores of *Pleurotus pulmonarius* were collected from a fruiting body. Two of the monokaryon strains (ss2 and ss5) each derived from a single basidiospore were established in the laboratory for analysis. Fungi were cultured on YMG medium (yeast extract, malt extract, and glucose) at 25°C. In order to extract their DNA, fungal mycelium was inoculated in 100 mL YMG liquid medium at 25°C for 3 days. DNA was extracted using cetyltrimethylammonium bromide (CTAB) and purified with chloroform, isopropanol, and phenol-chloroform. The genome of ss2 and ss5 was sequenced from the Pacbio RSII platform performed at Ramaciotti Centre for Genomics (Sydeny, Australia) using a PacBio SMRTBell Template Prep Kit 1.0 SPv3.

Fresh cultures of the *P. pulmonarius* strains were grown from mother cultures on YMG at 25°C for 7 days and then transferred to PDA (Potato Dextrose Agar) and LNM (Low Nutrient Medium) plates (5-cm diameter) for phenotypic characterization (in biological triplicates). After 13 days of growth, 30 adult *C. elegans* nematodes (strain N2) were added to the LNM plates and the number of paralyzed worms on each place was computed after 10 minutes.

### Genome sequencing and assembly

Long PacBio reads were sequenced from each *P. pulmonarius* strain with the Pacbio RSII platform. A total of ∼0.5M reads with a mean length of 9,419 bp were obtained from ss2, while ∼0.7M reads with a mean length of 8,422 bp were obtained from ss5; these corresponded to 122x genome coverage for ss2 and 142x genome coverage for ss5. The Pacbio reads were used to build two preliminary assemblies: one using Canu (v1.7) (Koren *et al*. 2017) and the other using Falcon (pbalign v0.0.2) (Chin *et al*. 2016). Canu was executed with parameters **genomeSize=40m useGrid=false maxThreads=8**, where the estimated genome size of 40Mbp was calculated as the average of the genome sizes of *Pleurotus eryngii* (44.61Mbp) and *Pleurotus ostreatus* (34.3-35.6Mbp). Falcon was run using an adapted version of the configuration file for fungal genomes assembling provided at: https://pb-falcon.readthedocs.io/en/latest/parameters.html, where **Genome_size** was set to 40Mb and memory-related options were changed to fit the requirements of PSC Bridges (Towns *et al*. 2014; Nystrom *et al*. 2015). Next, we used Sourmash (v2.0.0) (Titus Brown and Irber 2016) to identify and remove contaminants from each preliminary assembly. The decontaminated assemblies were subsequently polished using Quiver (genomicconsensus v2.3.2) (Chin *et al*. 2013) with the original raw PacBio reads, mapped to the decontaminated assembly using pbalign (v0.0.2), as input. Further information about the preliminary assemblies together with the results obtained from each of the abovementioned steps can be found in Supplementary Table S1.

The polished assemblies obtained with Canu were more accurate at the telomeric regions, while the Falcon assemblies were more contiguous. Hence, we merged the Canu and Falcon assemblies of each *P. pulmonarius* strain using Quickmerge (v0.3) (Chakraborty *et al*. 2016). Quickmerge uses the information of a donor assembly to fill in the gaps of a reference assembly, accordingly we tested both Canu as reference and Falcon as donor (Canu-Falcon) and *vice versa* (Falcon-Canu). Since the Canu-Falcon assemblies showed better contiguity and overall statistics (Supplementary Table S2), they were selected for further assembling steps. Nucmer (Mummer4) (Marçais *et al*. 2018) was used to detect redundant contigs in the Canu-Falcon assemblies (function maxmatch where the Canu-Falcon assembly was used both as reference and query). As a result, two small contigs (28,260 bp and 15,057 bp in size) were removed from the ss2 assembly, while only one contig (22,621 bp in size) was removed from the assembly of ss5. Finally, the contigs from the two assemblies were sorted by size and renamed.

### Genome annotation

The assemblies of *P. pulmonarius* were annotated with funannotate (v1.5.2) (Palmer and Stajich 2016) using the following steps and options. First, the genomes were softmasked using **funannotate mask**, calling RepeatMasker (Smit *et al*. 2013). The masked genomes were used as input for **funannotate train** (options: **--max_intronlen 2000 --stranded no**), together with RNA-seq data from *Pleurotus ostreatus* (strain PC9 (Alfaro *et al*. 2016), downloaded from the Joint Genome Institute (JGI) at https://mycocosm.jgi.doe.gov/PleosPC9_1/PleosPC9_1.info.html (Grigoriev *et al*. 2012)). During this step Trinity (Grabherr *et al*. 2013) and PASA (Haas *et al*. 2003) was run to generate preliminary gene models. Then, **funannotate predict** was used to generate consensus gene models from the preliminary gene models and known transcripts and proteins of *P. ostreatus* (use as evidence for gene prediction). **Funannotate predict** calls the following programs: AUGUSTUS (Stanke and Waack 2003), GeneMark (Borodovsky and McIninch 1993) and EVidenceModeler (Haas *et al*. 2008). In the final step of the annotation, **funannotate annotate** was run to add functional annotation to the final gene models including: InterPro and PFAM domains, GO ontology terms, fungal transcription factors, COGs, secondary metabolites (AntiSMASH (Medema *et al*. 2011) was run using **funnanotate remote**), CAZYmes, secreted proteins, proteases (MEROPS), and BUSCO groups.

### Genome analysis and comparison

General assembly statistics such as N50, length and number of scaffolds were computed directly from the fasta files using the Perl script count_fasta_residues.pl (https://github.com/SchwarzEM/ems_perl/blob/master/fasta/count_fasta_residues.pl). BUSCO completeness was assessed using BUSCO 3.0.1 (Simão *et al*. 2015) against the basidiomycota dataset basidiomycota_odb9. We used the protocol described in (Coghlan *et al*. 2018) to make repeat libraries and subsequently look for repetitive elements in each genome. We further identified and classified transposable elements using the one_code_to_find_them_all script (Bailly-Bechet *et al*. 2014). In addition, we searched for telomeric regions characterized by the telomeric repeating unit TTAGGG (Pérez *et al*. 2009). Raw PacBio reads were mapped to each genome using pbalign and SAMtools (Li *et al*. 2009) to compute local coverage.

In order to find structural differences between the two genomes, we used megablast (Morgulis *et al*. 2008) to precisely define the boundaries for regions of similarity and dissimilarity that were grossly visible in plots by Circos (Krzywinski *et al*. 2009) and Dgenies (relying on minimap2) (Cabanettes and Klopp 2018). Other comparisons between genomes were derived from the results of **funannotate compare** (Palmer and Stajich 2016).

### Data availability

The final assembled and annotated genomes of *P. pulmonarius* spores ss2 and ss5 were uploaded to NCBI and are available with accession codes: GCA_012980525.1 and GCA_012980535.1, respectively.

## Results

### Phenotypic differences between the two meiotic progeny of a *P. pulmonarius* mushroom

A wild *Pleurotus pulmonarius* mushroom (heterokaryon) was collected near Taipei, Taiwan and its basidiospores were collected and individually dissected using a dissecting microscope. Two monokaryotic strains derived from two basidiospores (referred as ss2 and ss5 strains from here on) were selected for genome sequencing because they exhibited prominently different phenotypes. For example, the growth of the two strains on Low Nutrient Medium (LNM) and on the rich Potato Dextrose Agar (PDA) were visibly dissimilar (Figure 1A). On nutrient-rich medium (PDA), ss2 developed a denser, but smaller mycelium colony compared to that of ss5. On low-nutrient medium, ss2 developed a smaller colony that exhibited similar hyphal density compared to that of ss5. In both conditions, the growth of ss5 was much faster than ss2. We further discovered that ss2 and ss5 exhibited differences in toxicity towards the nematode *Caenorhabditis elegans*. All nematodes in contact with the mycelium of ss5 were paralyzed after 10 minutes exposed to the fungal hyphae, which was comparable to previous studies of *Pleurotus* species (Thorn and Barron 1984; Lee *et al*. 2020). In contrast, the mycelium of ss2 showed a much weaker toxicity toward *C. elegans*, being able to paralyze only ∼40% of the nematode population tested (Figure 1B). In view of these results, we decided to further analyze the genetic differences of the two *P. pulmonarius* spores by sequencing their genomes.

**Figure 1.**
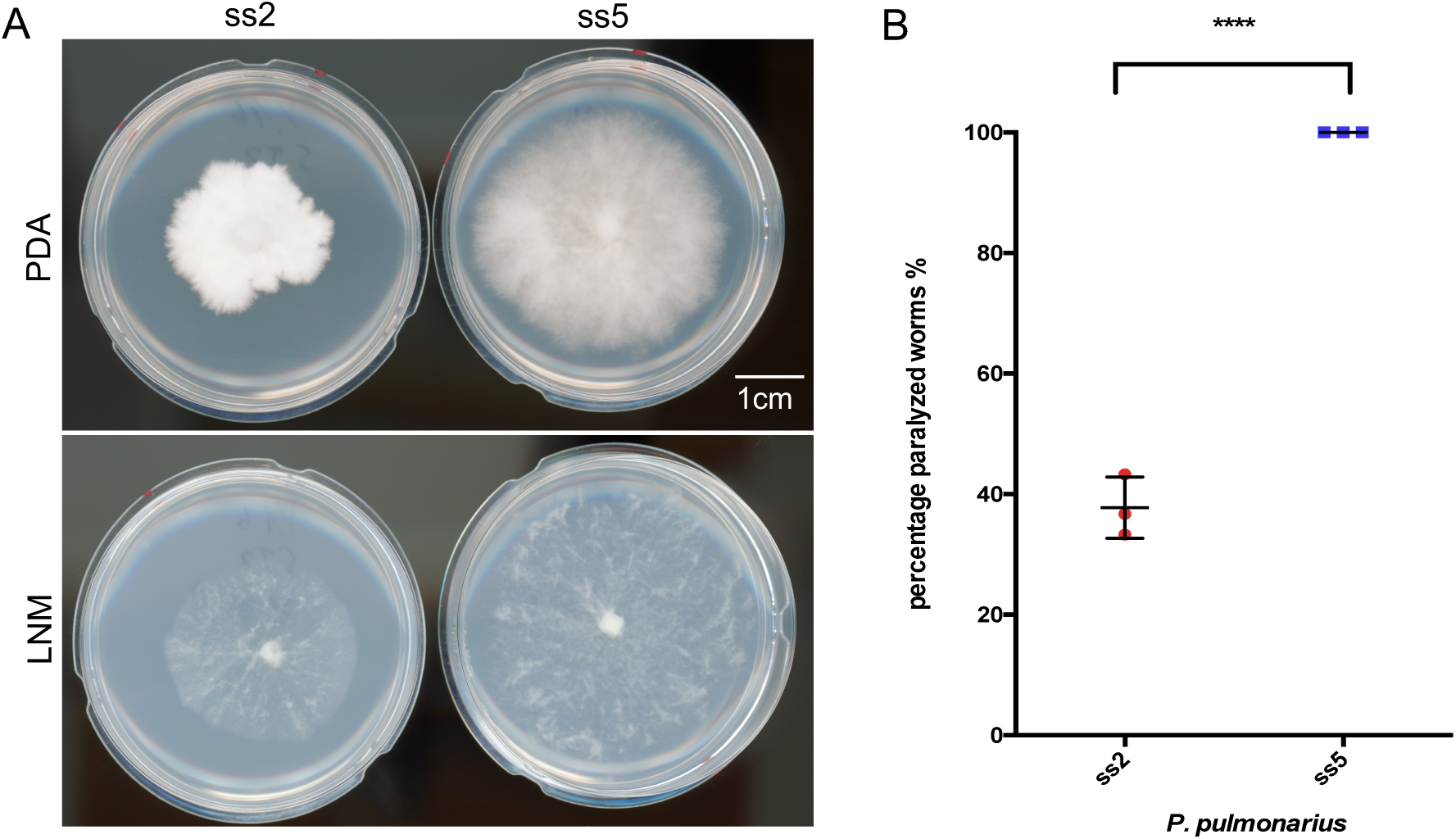
Phenotypic characterization of *P. pulmonarius* mycelia from ss2 and ss5. (A) Mycelia developed after 13 days of growth for each of the studied spores on Potato Dextrose Agar (PDA) and Low Nutrient Media (LNM). (B) Percentage of *C. elegans* (over a total of 30 nematodes) paralyzed after 10 minutes of contact with each *P. pulmonarius* strain.

### High quality genome assemblies of *P. pulmonarius*

Long PacBio reads were sequenced from both *P. pulmonarius* strains, ss2 and ss5, to build two high quality genome assemblies. The genome size of the assembled *P. pulmonarius* is ∼39.2 Mb for ss2 and ∼39.9 Mb for ss5 (Table 1). These sizes are consistent with genome sizes observed in other related *Pleurotus* species: for example, *P. ostreatus* with about 35 Mb (Riley *et al*. 2014; Qu *et al*. 2016; Wang *et al*. 2018) and *P. eryngii* with 49.9 Mb (Zhang *et al*. 2018). Both of our genomes are distributed in 23 scaffolds ranging in size from 5.55 Mb to 11 kb (in ss2) and 5.06 Mb to 21 kb (in ss5), and with N50 values of 3.2 Mb and 3.4 Mb (respectively), showing high contiguity. These N50 values are higher than those of four other available *Pleurotus* assemblies: *P. platypus* (N50 = 62 kb); *P. citrinopileatus* (N50 = 9.7 kb) (Li *et al*. 2018); *P. tuoliensis* (N50 = 1.17 Mb); and *P. eryngii* (N50 = 563 kb) (Zhang *et al*. 2018). Moreover, these N50 values are comparable to two of the best-quality *Pleurotus* genomes: *P. ostreatus* strain PC15 (N50 = 3.27 Mb) (Riley *et al*. 2014); and *P. ostreatus* strain CCMSSC00389 (N50 = 3.07 Mb) (Wang *et al*. 2018). We also computed the BUSCO completeness of both assemblies against the basidiomycota dataset; both assemblies showed high completeness (97.3% for ss2 and 96% for ss5). Overall, the *Pleurotus pulmonarius* assemblies presented here are among the most complete and contiguous of the *Pleurotus* species, only matched by the latest assemblies of *P. ostreatus* (strains PC15 (Riley *et al*. 2014) and CCMSSC00389 (Wang *et al*. 2018)).

**Table 1.** Genome features of *P. pulmonarius* strains ss2 and ss5.

| General features | *P. pulmonarius* ss2 | *P. pulmonarius* ss5 |
| --- | --- | --- |
| Genome size, bp | 39,235,597 | 39,866,212 |
| Number of scaffolds | 23 | 23 |
| GC content, % | 51 | 51.04 |
| Scaffold N50, bp | 3,175,356 | 3,374,720 |
| Scaffold N90, bp | 1,418,296 | 1,759,240 |
| Scaffold max size, bp | 5,554,685 | 5,063,380 |
| Scaffol min size, bp | 10,862 | 21,250 |
| Number of genes | 10,848 | 11,139 |
| Number of tRNA genes | 208 | 186 |
| BUSCO completeness, % | 97.3 | 96 |

The distribution and size of the scaffolds comprising the assembled genomes of *P. pulmonarius* is shown in Figure 2. First, we looked at the depth coverage at each genome position for both assemblies. While ss2 shows an average depth of 97.7 reads (Figure 2A), the average depth at each genome position of ss5 is only 86.36 reads (Figure 2B). High depth regions correspond mostly with the ends of the scaffolds for both genomes, whereas low depth is observed in some of the smaller scaffolds of both assemblies (scaffolds ss2.20 and ss2.22 for ss2, and ss5.18, ss5.19 and ss5.20 for ss5). Out of the 23 scaffolds of ss2, 6 scaffolds represent telomere-to-telomere complete chromosomes, while another 14 scaffolds show only one telomere at one end (Figure 2A). The genome of ss5 is distributed in 8 whole chromosomes and 8 half-chromosomes (Figure 2B). Additionally, we observed telomeric repeats in several of the small scaffolds; this suggests that they may correspond to the other ends of some half-chromosomes, but failed to be assembled into these larger contigs due to their repetitive nature. These results are in line with the total number of chromosomes estimated in other *Pleurotus* species such as *P. ostreatus* (11 chromosomes) (Larraya *et al*. 2000; Pérez *et al*. 2009) and *P. tuoliensis* (12 chromosomes) (Gao *et al*. 2018). Next, we generated a custom-built repeat library for each genome to study the distribution of transposable elements (TE). About 7.3% (6.4%) of the genome of *P. pulmonarius* ss2 (ss5) consists of transposable elements (Table 2). The repetitive sequences in of our *P. pulmonarius* genome assemblies mainly contain class I elements, retrotransposons, accounting for ∼90% and ∼80% of the repetitive genome fraction and about 78% and 85% of the total repeats in ss2 and ss5, respectively. TE density and distribution can be also observed in Figure 2; these are highly variable, as has been previously reported for other *Pleurotus* species (Castanera *et al*. 2016). We observed clusters of TEs in both genomes mostly aligning with regions of relative low gene density (Figure 2). However, we did not observe clear evidence of centromeres, which are often characterized by clusters of transposable elements in fungi (Stajich *et al*. 2010). Detailed information about each of the classified TE identified in ss2 and ss5 can be found in Supplementary Tables TS3 and TS4, respectively. Gene density was computed for each assembly (Figure 2) in sliding windows of 100Kb. A total of 10,848 genes were predicted in ss2 using funannotate (Palmer and Stajich 2016) and 11,139 were found in ss5 genome. These values are within the range of close *Pleurotus* species, such as *P. ostreatus* (11,603 genes in PC15 and 12,206 in PC9 (Castanera *et al*. 2016)) and *P. eryngii* (15,960 genes in strain ATCC 90797 (Gao *et al*. 2018)) as described in jgi (Grigoriev *et al*. 2012). On average about 28 genes per 100-kb sliding window were annotated in both ss2 and ss5, however genes are not evenly distributed across the genomes presenting areas of low and high gene density for most scaffolds (Figure 2). Moreover, few genes were predicted in the smaller scaffolds of both genomes, once again suggesting that these scaffolds may correspond to misplaced telomeric or centromeric regions of chromosomes. The predicted function of genes in both genomes, cataloged using the cluster of orthologous groups (COGs) database, can be found in Figure 2C. As hinted by the difference in total number of annotated genes between the genomes, ss5 contains more genes than ss2 for most functional categories. However, significant differences in the overall distribution of genes by function between the two genomes were not observed (Figure 2C).

**Table 2.** Classification of transposable elements identified in *P. pulmonarius* monokaryon strains ss2 and ss5.

| Family | *P. pulmonarius* ss2 |  |  | *P. pulmonarius* ss5 |  |  |
| --- | --- | --- | --- | --- | --- | --- |
|  | Fragments | Copies | Total Bp | Fragments | Copies | Total Bp |
| DNA/CMC-EnSpm | 142 | 82 | 88249 | 126 | 47 | 93887 |
| DNA/Crypton | 5 | 4 | 25088 | 0 | 0 | 0 |
| DNA/Crypton-V | 0 | 0 | 0 | 54 | 51 | 316697 |
| DNA/Dada | 0 | 0 | 0 | 12 | 10 | 9340 |
| DNA/hAT-Ac | 10 | 7 | 6624 | 4 | 1 | 786 |
| DNA/Maverick | 152 | 52 | 41933 | 39 | 20 | 51514 |
| DNA/MULE-MuDR | 77 | 17 | 41283 | 26 | 3 | 1870 |
| DNA/PIF-Harbinger | 19 | 9 | 8138 | 13 | 3 | 1103 |
| DNA/RC | 95 | 59 | 78957 | 33 | 22 | 44417 |
| DNA/TcMar-Ant1 | 11 | 7 | 5873 | 3 | 1 | 981 |
| DNA/TcMar-Fot1 | 10 | 7 | 9998 | 8 | 6 | 9088 |
| DNA/TcMar-Pogo | 2 | 2 | 1284 | 3 | 1 | 540 |
| DNA/TcMar-Tc1 | 3 | 3 | 2578 | 0 | 0 | 0 |
| Other DNA | 5 | 3 | 5289 | 37 | 9 | 7742 |
| LINE/I | 1 | 1 | 414 | 1 | 0 | 0 |
| LINE/L1 | 107 | 5 | 11665 | 19 | 11 | 23464 |
| LINE/L1-Tx1 | 3 | 1 | 2451 | 21 | 8 | 23115 |
| LINE/L2 | 157 | 102 | 651829 | 101 | 93 | 184053 |
| LINE/R1 | 64 | 41 | 21126 | 0 | 0 | 0 |
| LINE/Tad1 | 48 | 37 | 56788 | 20 | 14 | 28805 |
| Other LINE | 5 | 1 | 834 | 0 | 0 | 0 |
| LTR/Caulimovirus | 0 | 0 | 0 | 55 | 21 | 23920 |
| LTR/Copia | 227 | 134 | 299643 | 425 | 167 | 292397 |
| LTR/Gypsy | 947 | 525 | 1413722 | 1117 | 641 | 1380124 |
| LTR/NGaro | 128 | 33 | 87204 | 60 | 36 | 61347 |
| LTR/Pao | 0 | 0 | 0 | 12 | 4 | 4632 |
| Other LTR | 16 | 16 | 5979 | 18 | 18 | 6256 |
| **Total Repeats** | **2218** | **1132** | **2860970** | **2189** | **1169** | **2559822** |

**Figure 2.**
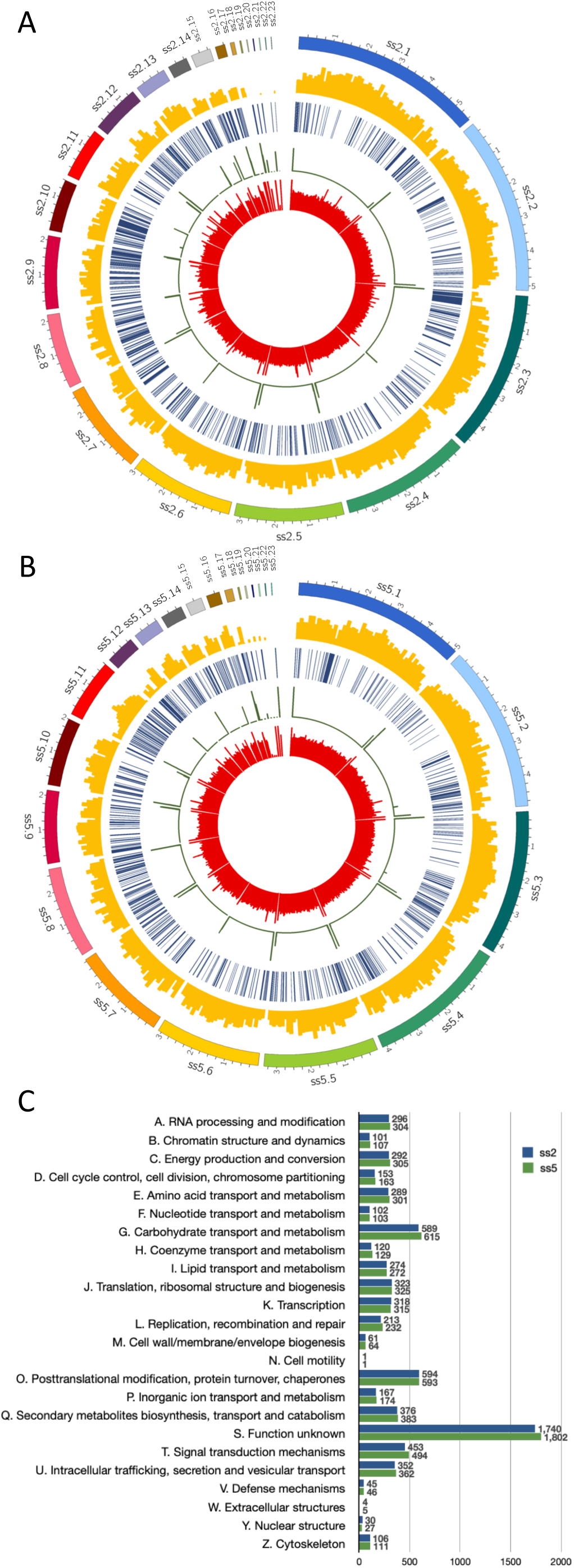
Genome architecture of *P. pulmonarius*. Tracks (outer to inner) represent the distribution of genomic features: 1) positions (in Mb) of the assembled scaffolds, with numbers indicating the order of scaffolds by size; 2) gene density (along 100-kb sliding windows, ranging between 0-50 genes); 3) distribution of transposable elements along the genome; 4) telomere repeat frequency (along 10kb sliding window, ranging between 10-30 repeats); and 5) Mean base pair depth coverage (along a 10 kb sliding windows, ranging between 0-150 depth) for strains ss2 (A) and ss5 (B). (C) Predicted function of genes in each *P. pulmonarius* assembly catalogued using the cluster of orthologous groups (COGs) database.

### Genomic differences between *P. pulmonarius* strains

Using our high-quality assemblies of *P. pulmonarius*, we conducted a preliminary analysis of the main differences between the genomes of ss2 and ss5. As can be observed in the dot-plot of Figure 3A, our assemblies aligned almost perfectly to each other, only showing lower identity at the ends of the scaffolds and a few regions of low similarity. Moreover, we observed that the end of scaffold10 of ss2 and the beginning of scaffold9 of ss5 do not correspond to any of the major scaffolds of the opposite assembly (Figure 3A). A closer look to the similarities between both assemblies can be found in Figure 3B, showing high similarity regions (>95%) between the most relevant scaffolds of each genome (>500 kb in size). In this comparison, the missing beginning of scaffold 9 in ss5 and end of scaffold 10 in ss2 become even more apparent. We observed some possible chromosomal translocation in scaffold 2 of ss5 consisting of the majority of scaffold 1 and a piece of scaffold 10 from ss2; however, due to the presence of many transposable elements in the genome, we cannot rule out the possibility that this is a result of misassembly. Further experimental analysis would be necessary to confirm this phenomenon. Finally, we looked for genes that are present only in one of the genomes. A prominent cluster of unique genes at the beginning of scaffold 9 in ss5 contained 159 genes that are missing in ss2 (Supplementary table TS6). Similarly, we also observed that ss2 harbored 66 genes on scaffold 10 that were not present in ss5 (Supplementary table TS5).

**Figure 3.**
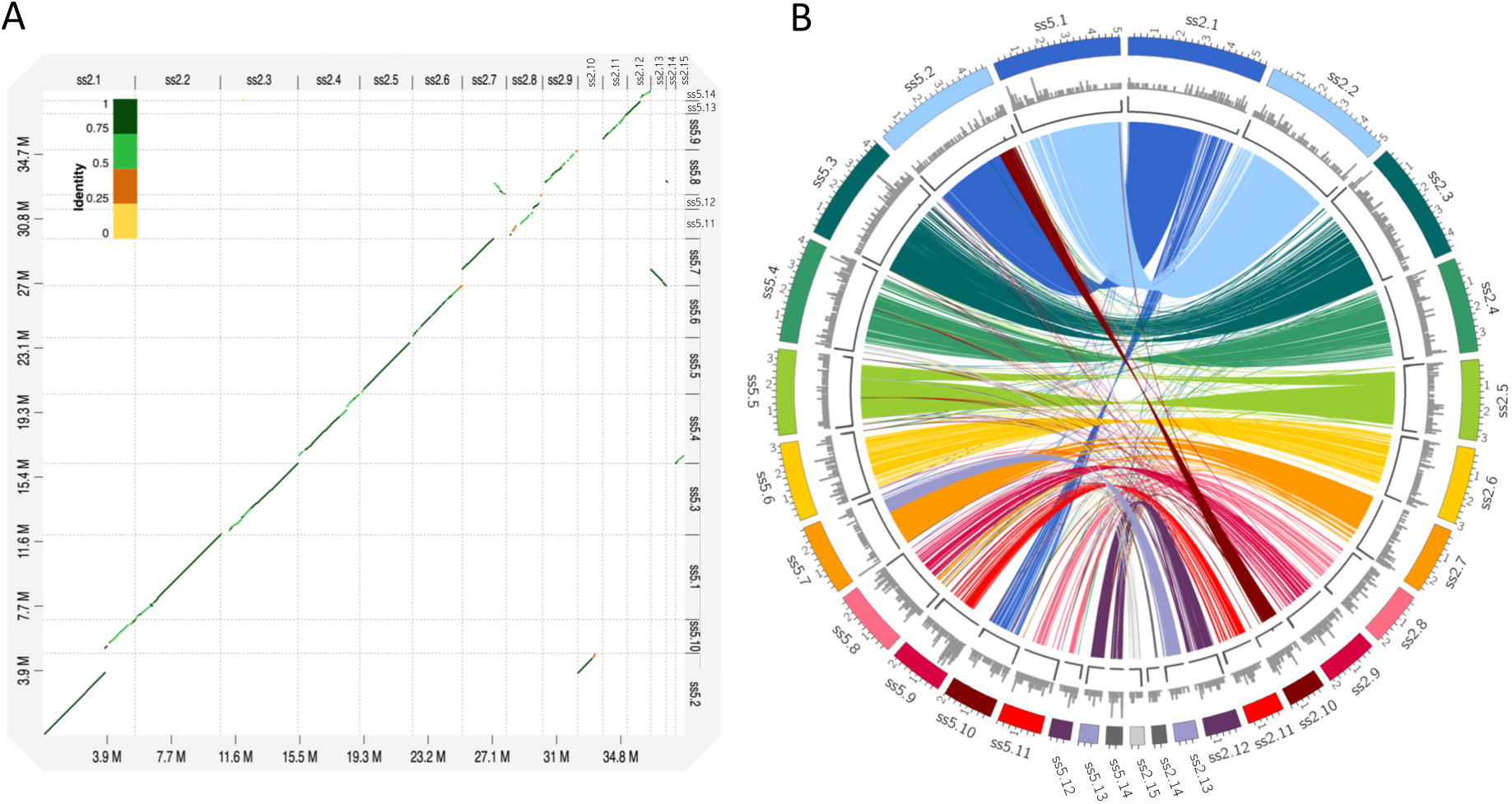
Comparison between the assembled genomes of *P. pulmonarius*, strains ss2 and ss5. (A) Dot-plot alignment of ss5 (query) against ss2 (reference). (B) Correspondence between regions of high similarity (>95%, length > 10Kb) among the most relevant scaffolds (>500 kb in size) of the two assembled genomes. Tracks (outer to inner) represent the distribution of genomic features in each assembly: 1) positions (in Mb) of the assembled scaffolds; 2) number of genes unique in each assembly (along 100-kb sliding windows, ranging between 0-35 genes); and 3) telomere repeat frequency (along 10-kb sliding window, ranging between 10-30 repeats).

## Conclusions

In this study we conducted PacBio genome sequencing for two spores harvested from a dikaryotic strain of *Pleurotus pulmonarius* and assembled two high quality genomes. We observed significant phenotypic differences in terms of growth and toxicity against nematodes between the two monokaryotic strains. Motivated by these results, we conducted genomic analyses of these two *P. pulmonarius* strains. Their two assemblies are the first available genome resources for *P. pulmonarius*, and their quality places them among the best *Pleurotus* genomes. Comparison of these two assemblies showed dissimilar regions that may contribute to the phenotypic differences of these two strains. The *de novo* assemblies presented here will be crucial genomic resources for studying the biology of *P. pulmonarius*; they will also enable comparative genomic studies in *Pleurotus* mushrooms, which contains several species of most cultivated mushrooms around the globe.

## Acknowledgments

The authors thank Ting-Fang Wang for comments on this manuscript. Work here was supported in part by startup and research allocations from NSF XSEDE (TG-MCB180039 and TG-MCB190010) to EMS and Academia Sinica Career Development Award AS-CDA-106-L03 to YPH.

## Supplemental information

**Table S1.** Features of the preliminary assemblies of *P. pulmonarius* ss2 and ss5 obtained with Canu, Falcon, Sourmash (decontamination) and Quiver (polishing).

|  | Features | Canu | Canu +<br>Sourmash | Canu + Sourmash<br>+ Quiver | Falcon | Falcon +<br>Sourmash | Falcon + Sourmash<br>+ Quiver |
| --- | --- | --- | --- | --- | --- | --- | --- |
| *Pleurotus pulmonarius*<br>ss2 | Total nt | 40,950,889 | 40,774,743 | 40,778,513 | 40,527,552 | 40,383,578 | 40,430,866 |
|  | Scaffolds | 54 | 50 | 50 | 39 | 34 | 34 |
|  | ACGT nt | 40,950,889 | 40,774,743 |  | 40,527,552 | 40,383,578 | 40,430,866 |
|  | % GC | 50.93 | 50.98 | 50.99 | 50.93 | 50.97 | 51.01 |
|  | Scaffold N50 nt | 3,012,406.00 | 3,012,406.00 | 3,012,621.00 | 2,238,199.00 | 2,638,868.00 | 2,641,788.00 |
|  | Scaffold N90 nt | 974,086.00 | 974,086.00 | 974,184.00 | 939,585.00 | 939,585.00 | 940,770.00 |
|  | Scaf. max. nt | 5,139,728 | 5,139,728 | 5,140,519 | 5,135,074 | 5,135,074 | 5,140,517 |
|  | Scaf. min. nt | 2,843 | 2,843 | 2,841 | 10,574 | 10,574 | 10,862 |
| *Pleurotus pulmonarius*<br>ss5 | Total nt | 40,631,668 | 40,449,693 | 40,450,376 | 39,969,076 | 39,945,311 | 39,961,457 |
|  | Scaffolds | 59 | 55 | 55 | 30 | 29 | 29 |
|  | ACGT nt | 40,631,668 | 40,449,693 | 40,450,376 | 39,969,076 | 39,945,311 | 39,961,457 |
|  | % GC | 50.94 | 50.97 | 50.97 | 51.04 | 51.04 | 51.05 |
|  | Scaffold N50 nt | 3,030,066.00 | 3,030,066.00 | 3,028,859.00 | 3,100,931.00 | 3,100,931.00 | 3,102,052.00 |
|  | Scaffold N90 nt | 773,592.00 | 773,592.00 | 773,543.00 | 866,022.00 | 866,022.00 | 866,283.00 |
|  | Scaf. max. nt | 5,063,046 | 5,063,046 | 5,063,242 | 5,061,483 | 5,061,483 | 5,063,380 |
|  | Scaf. min. nt | 1,546 | 1,546 | 1,549 | 14,558 | 14,558 | 14,587 |

**Table S2.**
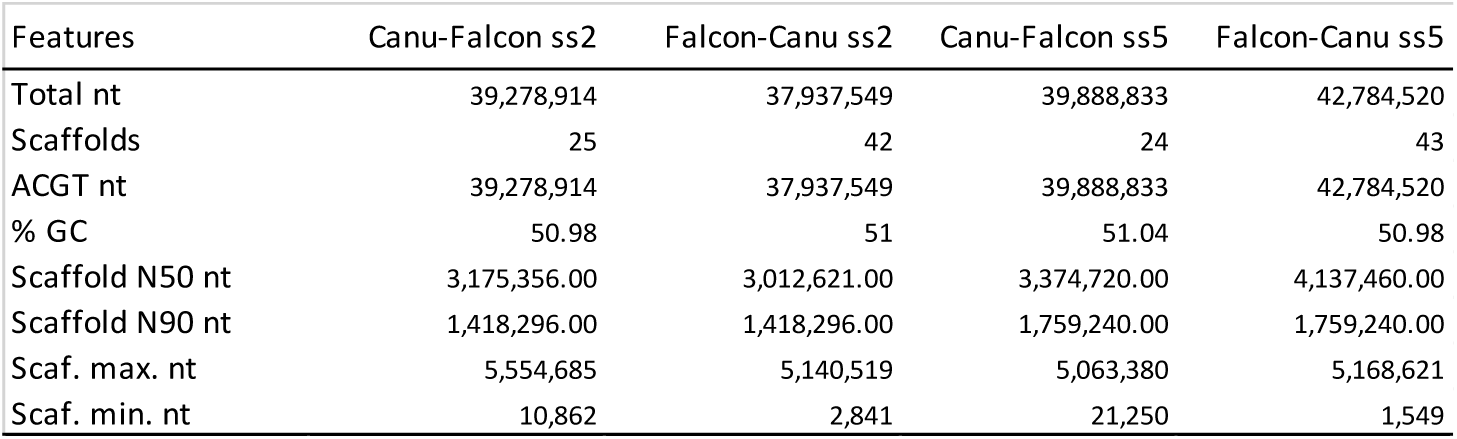
Features of the merged assemblies of *P. pulmonarius* ss2 and ss5 constructed with Quickmerge using the Canu assembly as reference and the Falcon assembly as donor (Canu-Falcon) and *vice versa* (Falcon-Canu).

| Features | Canu-Falcon ss2 | Falcon-Canu ss2 | Canu-Falcon ss5 | Falcon-Canu ss5 |
| --- | --- | --- | --- | --- |
| Total nt | 39,278,914 | 37,937,549 | 39,888,833 | 42,784,520 |
| Scaffolds | 25 | 42 | 24 | 43 |
| ACGT nt | 39,278,914 | 37,937,549 | 39,888,833 | 42,784,520 |
| % GC | 50.98 | 51 | 51.04 | 50.98 |
| Scaffold N50 nt | 3,175,356.00 | 3,012,621.00 | 3,374,720.00 | 4,137,460.00 |
| Scaffold N90 nt | 1,418,296.00 | 1,418,296.00 | 1,759,240.00 | 1,759,240.00 |
| Scaf. max. nt | 5,554,685 | 5,140,519 | 5,063,380 | 5,168,621 |
| Scaf. min. nt | 10,862 | 2,841 | 21,250 | 1,549 |

**Table S3.** Detected TE elements in *P. pulmonarius* ss2.

| Element | Classification | Size | Fragments | Repeats | Total Bp |
| --- | --- | --- | --- | --- | --- |
| ss2_TE_1 | DNA/TcMar-Tc1 | 984 | 1 | 1 | 984 |
| ss2_TE_2 | DNA/TcMar-Tc1 | 972 | 2 | 2 | 1594 |
| ss2_TE_3 | DNA/CMC-EnSpm | 1692 | 3 | 1 | 1692 |
| ss2_TE_4 | DNA/CMC-EnSpm | 1119 | 9 | 3 | 2311 |
| ss2_TE_5 | DNA/CMC-EnSpm | 612 | 1 | 1 | 612 |
| ss2_TE_6 | DNA/CMC-EnSpm | 1692 | 4 | 1 | 1692 |
| ss2_TE_7 | DNA/CMC-EnSpm | 407 | 6 | 6 | 2418 |
| ss2_TE_8 | DNA/CMC-EnSpm | 573 | 2 | 0 | 0 |
| ss2_TE_9 | DNA/CMC-EnSpm | 2393 | 23 | 10 | 15721 |
| ss2_TE_10 | DNA/CMC-EnSpm | 609 | 2 | 2 | 1218 |
| ss2_TE_11 | DNA/CMC-EnSpm | 837 | 2 | 2 | 1674 |
| ss2_TE_12 | DNA/CMC-EnSpm | 420 | 1 | 1 | 420 |
| ss2_TE_13 | DNA/CMC-EnSpm | 908 | 22 | 16 | 5157 |
| ss2_TE_14 | DNA/CMC-EnSpm | 903 | 3 | 1 | 902 |
| ss2_TE_15 | DNA/CMC-EnSpm | 1289 | 13 | 10 | 8327 |
| ss2_TE_16 | DNA/CMC-EnSpm | 1119 | 1 | 1 | 1117 |
| ss2_TE_17 | DNA/CMC-EnSpm | 1692 | 3 | 1 | 892 |
| ss2_TE_18 | DNA/CMC-EnSpm | 882 | 1 | 1 | 882 |
| ss2_TE_19 | DNA/CMC-EnSpm | 666 | 5 | 5 | 3330 |
| ss2_TE_20 | DNA/CMC-EnSpm | 8078 | 19 | 8 | 33608 |
| ss2_TE_21 | DNA/CMC-EnSpm | 900 | 3 | 3 | 2700 |
| ss2_TE_22 | DNA/CMC-EnSpm | 441 | 19 | 9 | 3576 |
| ss2_TE_23 | DNA/RC | 870 | 7 | 7 | 6090 |
| ss2_TE_24 | DNA/RC | 1233 | 4 | 4 | 2632 |
| ss2_TE_25 | DNA/RC | 4155 | 1 | 0 | 0 |
| ss2_TE_26 | DNA/RC | 4248 | 3 | 1 | 4248 |
| ss2_TE_27 | DNA/RC | 4164 | 2 | 2 | 8328 |
| ss2_TE_28 | DNA/RC | 4155 | 1 | 1 | 4152 |
| ss2_TE_29 | DNA/RC | 3943 | 2 | 1 | 3943 |
| ss2_TE_30 | DNA/RC | 634 | 5 | 3 | 1080 |
| ss2_TE_31 | DNA/RC | 1713 | 1 | 1 | 1713 |
| ss2_TE_32 | DNA/RC | 1527 | 1 | 1 | 1527 |
| ss2_TE_33 | DNA/RC | 1477 | 6 | 6 | 8862 |
| ss2_TE_34 | DNA/RC | 1533 | 4 | 3 | 4599 |
| ss2_TE_35 | DNA/RC | 1551 | 1 | 1 | 1551 |
| ss2_TE_36 | DNA/RC | 1437 | 1 | 1 | 1437 |
| ss2_TE_37 | DNA/RC | 720 | 3 | 1 | 720 |
| ss2_TE_38 | DNA/RC | 1492 | 1 | 1 | 1492 |
| ss2_TE_39 | DNA/RC | 1461 | 1 | 1 | 1461 |
| ss2_TE_40 | DNA/RC | 4179 | 2 | 2 | 4846 |
| ss2_TE_41 | DNA/RC | 906 | 5 | 3 | 2718 |
| ss2_TE_42 | DNA/RC | 4318 | 31 | 15 | 6239 |
| ss2_TE_43 | DNA/RC | 4155 | 1 | 1 | 4155 |
| ss2_TE_44 | DNA/RC | 4236 | 6 | 1 | 4236 |
| ss2_TE_45 | DNA/RC | 1437 | 1 | 1 | 1437 |
| ss2_TE_46 | DNA/RC | 1491 | 5 | 1 | 1491 |
| ss2_TE_47 | DNA/TcMar-Fot1 | 894 | 2 | 1 | 894 |
| ss2_TE_48 | DNA/TcMar-Fot1 | 1286 | 1 | 1 | 1286 |
| ss2_TE_49 | DNA/TcMar-Fot1 | 1722 | 1 | 1 | 1706 |
| ss2_TE_50 | DNA/TcMar-Fot1 | 1342 | 1 | 1 | 1342 |
| ss2_TE_51 | DNA/TcMar-Fot1 | 1722 | 3 | 1 | 1722 |
| ss2_TE_52 | DNA/TcMar-Fot1 | 1316 | 1 | 1 | 1316 |
| ss2_TE_53 | DNA/TcMar-Fot1 | 1732 | 1 | 1 | 1732 |
| ss2_TE_54 | DNA/TcMar-Ant1 | 1016 | 1 | 1 | 835 |
| ss2_TE_55 | DNA/TcMar-Ant1 | 1016 | 4 | 2 | 2032 |
| ss2_TE_56 | DNA/TcMar-Ant1 | 916 | 1 | 1 | 916 |
| ss2_TE_57 | DNA/TcMar-Ant1 | 1016 | 1 | 0 | 0 |
| ss2_TE_58 | DNA/TcMar-Ant1 | 657 | 1 | 1 | 657 |
| ss2_TE_59 | DNA/TcMar-Ant1 | 573 | 2 | 1 | 573 |
| ss2_TE_60 | DNA/TcMar-Ant1 | 860 | 1 | 1 | 860 |
| ss2_TE_61 | DNA/hAT-Ac | 768 | 3 | 3 | 2304 |
| ss2_TE_62 | DNA/hAT-Ac | 606 | 2 | 2 | 1212 |
| ss2_TE_63 | DNA/hAT-Ac | 1737 | 1 | 1 | 1737 |
| ss2_TE_64 | DNA/hAT-Ac | 1371 | 4 | 1 | 1371 |
| ss2_TE_65 | DNA/MULE-MuDR | 718 | 10 | 2 | 634 |
| ss2_TE_66 | DNA/MULE-MuDR | 3878 | 67 | 15 | 40649 |
| ss2_TE_67 | DNA | 1647 | 3 | 1 | 1647 |
| ss2_TE_68 | DNA | 1821 | 2 | 2 | 3642 |
| ss2_TE_69 | DNA/PIF-Harbinger | 1705 | 19 | 9 | 8138 |
| ss2_TE_70 | DNA/MuLE-MuDR | 537 | 1 | 1 | 537 |
| ss2_TE_71 | DNA/TcMar-Pogo | 744 | 1 | 1 | 744 |
| ss2_TE_72 | DNA/TcMar-Pogo | 540 | 1 | 1 | 540 |
| ss2_TE_73 | DNA/Crypton | 6272 | 5 | 4 | 25088 |
| ss2_TE_74 | DNA/Maverick | 5530 | 152 | 52 | 41933 |
| ss2_TE_75 | LINE/Tad1 | 2756 | 1 | 1 | 3595 |
| ss2_TE_76 | LINE/Tad1 | 2403 | 1 | 1 | 3595 |
| ss2_TE_77 | LINE/Tad1 | 1020 | 2 | 2 | 2040 |
| ss2_TE_78 | LINE/Tad1 | 2756 | 1 | 0 | 0 |
| ss2_TE_79 | LINE/Tad1 | 3396 | 2 | 2 | 6792 |
| ss2_TE_80 | LINE/Tad1 | 3702 | 3 | 1 | 3702 |
| ss2_TE_81 | LINE/Tad1 | 4261 | 30 | 23 | 17603 |
| ss2_TE_82 | LINE/Tad1 | 1341 | 1 | 1 | 1341 |
| ss2_TE_83 | LINE/Tad1 | 3666 | 5 | 4 | 13407 |
| ss2_TE_84 | LINE/Tad1 | 2403 | 2 | 2 | 4713 |
| ss2_TE_85 | LINE/L1 | 1567 | 99 | 3 | 3320 |
| ss2_TE_86 | LINE/L1 | 4209 | 8 | 2 | 8345 |
| ss2_TE_87 | LINE/R1 | 2539 | 64 | 41 | 21126 |
| ss2_TE_88 | LINE/L1-Tx1 | 2451 | 3 | 1 | 2451 |
| ss2_TE_89 | LINE/L2 | 13558 | 157 | 102 | 651829 |
| ss2_TE_90 | LINE/I | 414 | 1 | 1 | 414 |
| ss2_TE_91 | LINE | 834 | 5 | 1 | 834 |
| ss2_TE_92 | LTR/Copia | 5898 | 14 | 12 | 32796 |
| ss2_TE_93 | LTR/Copia | 834 | 1 | 1 | 834 |
| ss2_TE_94 | LTR/Copia | 6596 | 2 | 2 | 10444 |
| ss2_TE_95 | LTR/Copia | 5893 | 9 | 9 | 17789 |
| ss2_TE_96 | LTR/Copia | 5775 | 15 | 13 | 22042 |
| ss2_TE_97 | LTR/Copia | 3270 | 1 | 1 | 657 |
| ss2_TE_98 | LTR/Copia | 6152 | 5 | 5 | 7086 |
| ss2_TE_99 | LTR/Copia | 5891 | 12 | 11 | 15697 |
| ss2_TE_100 | LTR/Copia | 5854 | 6 | 6 | 9990 |
| ss2_TE_101 | LTR/Copia | 14730 | 88 | 25 | 54212 |
| ss2_TE_102 | LTR/Copia | 3846 | 1 | 1 | 3846 |
| ss2_TE_103 | LTR/Copia | 5900 | 19 | 15 | 53365 |
| ss2_TE_104 | LTR/Copia | 3270 | 5 | 2 | 3479 |
| ss2_TE_105 | LTR/Copia | 1534 | 1 | 1 | 1534 |
| ss2_TE_106 | LTR/Copia | 6614 | 1 | 1 | 6614 |
| ss2_TE_107 | LTR/Copia | 5032 | 2 | 2 | 10064 |
| ss2_TE_108 | LTR/Copia | 1426 | 5 | 5 | 2271 |
| ss2_TE_109 | LTR/Copia | 6413 | 9 | 9 | 12275 |
| ss2_TE_110 | LTR/Copia | 2493 | 1 | 1 | 357 |
| ss2_TE_111 | LTR/Copia | 6600 | 3 | 2 | 9356 |
| ss2_TE_112 | LTR/Copia | 5488 | 26 | 9 | 24491 |
| ss2_TE_113 | LTR/Copia | 444 | 1 | 1 | 444 |
| ss2_TE_114 | LTR | 543 | 16 | 16 | 5979 |
| ss2_TE_115 | LTR/Ngaro | 4603 | 3 | 1 | 3136 |
| ss2_TE_116 | LTR/Ngaro | 4619 | 5 | 4 | 9587 |
| ss2_TE_117 | LTR/Ngaro | 2685 | 1 | 1 | 2685 |
| ss2_TE_118 | LTR/Ngaro | 4635 | 4 | 3 | 10169 |
| ss2_TE_119 | LTR/Ngaro | 4578 | 2 | 2 | 4746 |
| ss2_TE_120 | LTR/Ngaro | 4770 | 17 | 2 | 5022 |
| ss2_TE_121 | LTR/Ngaro | 5030 | 5 | 1 | 5030 |
| ss2_TE_122 | LTR/Ngaro | 4603 | 4 | 2 | 4772 |
| ss2_TE_123 | LTR/Ngaro | 4781 | 28 | 8 | 14521 |
| ss2_TE_124 | LTR/Ngaro | 4608 | 4 | 3 | 4953 |
| ss2_TE_125 | LTR/Ngaro | 13708 | 55 | 6 | 22583 |
| ss2_TE_126 | LTR/Gypsy | 10379 | 2 | 1 | 10379 |
| ss2_TE_127 | LTR/Gypsy | 8281 | 9 | 1 | 8281 |
| ss2_TE_128 | LTR/Gypsy | 4864 | 2 | 1 | 4864 |
| ss2_TE_129 | LTR/Gypsy | 6698 | 3 | 3 | 7812 |
| ss2_TE_130 | LTR/Gypsy | 6707 | 1 | 1 | 6707 |
| ss2_TE_131 | LTR/Gypsy | 5688 | 1 | 1 | 3276 |
| ss2_TE_132 | LTR/Gypsy | 7918 | 37 | 7 | 17561 |
| ss2_TE_133 | LTR/Gypsy | 5654 | 4 | 3 | 15882 |
| ss2_TE_134 | LTR/Gypsy | 510 | 1 | 1 | 510 |
| ss2_TE_135 | LTR/Gypsy | 9909 | 1 | 1 | 9909 |
| ss2_TE_136 | LTR/Gypsy | 6704 | 4 | 4 | 8109 |
| ss2_TE_137 | LTR/Gypsy | 8281 | 1 | 1 | 5451 |
| ss2_TE_138 | LTR/Gypsy | 6704 | 3 | 3 | 12703 |
| ss2_TE_139 | LTR/Gypsy | 2001 | 2 | 2 | 4281 |
| ss2_TE_140 | LTR/Gypsy | 4714 | 67 | 28 | 52306 |
| ss2_TE_141 | LTR/Gypsy | 1737 | 34 | 11 | 10445 |
| ss2_TE_142 | LTR/Gypsy | 2453 | 1 | 0 | 0 |
| ss2_TE_143 | LTR/Gypsy | 10333 | 10 | 5 | 10993 |
| ss2_TE_144 | LTR/Gypsy | 6694 | 1 | 1 | 6694 |
| ss2_TE_145 | LTR/Gypsy | 9887 | 1 | 1 | 9887 |
| ss2_TE_146 | LTR/Gypsy | 7983 | 10 | 9 | 15234 |
| ss2_TE_147 | LTR/Gypsy | 6690 | 2 | 1 | 6690 |
| ss2_TE_148 | LTR/Gypsy | 7956 | 15 | 9 | 18841 |
| ss2_TE_149 | LTR/Gypsy | 6700 | 2 | 1 | 6700 |
| ss2_TE_150 | LTR/Gypsy | 8826 | 1 | 1 | 8826 |
| ss2_TE_151 | LTR/Gypsy | 2613 | 20 | 19 | 17751 |
| ss2_TE_152 | LTR/Gypsy | 6715 | 1 | 1 | 6715 |
| ss2_TE_153 | LTR/Gypsy | 6709 | 3 | 3 | 7906 |
| ss2_TE_154 | LTR/Gypsy | 9428 | 22 | 16 | 33648 |
| ss2_TE_155 | LTR/Gypsy | 6691 | 2 | 1 | 6691 |
| ss2_TE_156 | LTR/Gypsy | 6683 | 1 | 1 | 6683 |
| ss2_TE_157 | LTR/Gypsy | 6711 | 1 | 1 | 6711 |
| ss2_TE_158 | LTR/Gypsy | 6711 | 2 | 1 | 6711 |
| ss2_TE_159 | LTR/Gypsy | 6709 | 2 | 2 | 6828 |
| ss2_TE_160 | LTR/Gypsy | 6674 | 1 | 1 | 6674 |
| ss2_TE_161 | LTR/Gypsy | 3270 | 6 | 6 | 1339 |
| ss2_TE_162 | LTR/Gypsy | 6709 | 2 | 2 | 7314 |
| ss2_TE_163 | LTR/Gypsy | 6699 | 3 | 3 | 16468 |
| ss2_TE_164 | LTR/Gypsy | 14332 | 20 | 18 | 27882 |
| ss2_TE_165 | LTR/Gypsy | 13930 | 13 | 5 | 33719 |
| ss2_TE_166 | LTR/Gypsy | 10267 | 2 | 2 | 10860 |
| ss2_TE_167 | LTR/Gypsy | 688 | 9 | 7 | 981 |
| ss2_TE_168 | LTR/Gypsy | 5654 | 4 | 2 | 5829 |
| ss2_TE_169 | LTR/Gypsy | 2830 | 6 | 5 | 8240 |
| ss2_TE_170 | LTR/Gypsy | 6719 | 2 | 2 | 11256 |
| ss2_TE_171 | LTR/Gypsy | 4369 | 6 | 3 | 4605 |
| ss2_TE_172 | LTR/Gypsy | 7300 | 3 | 3 | 11489 |
| ss2_TE_173 | LTR/Gypsy | 6756 | 42 | 5 | 32783 |
| ss2_TE_174 | LTR/Gypsy | 8331 | 1 | 1 | 8331 |
| ss2_TE_175 | LTR/Gypsy | 6115 | 4 | 4 | 7955 |
| ss2_TE_176 | LTR/Gypsy | 4912 | 27 | 6 | 10533 |
| ss2_TE_177 | LTR/Gypsy | 1194 | 5 | 4 | 4524 |
| ss2_TE_178 | LTR/Gypsy | 8007 | 1 | 1 | 8007 |
| ss2_TE_179 | LTR/Gypsy | 6708 | 1 | 1 | 6708 |
| ss2_TE_180 | LTR/Gypsy | 5668 | 5 | 4 | 11651 |
| ss2_TE_181 | LTR/Gypsy | 2685 | 2 | 2 | 5370 |
| ss2_TE_182 | LTR/Gypsy | 6440 | 3 | 3 | 1815 |
| ss2_TE_183 | LTR/Gypsy | 6690 | 2 | 2 | 13380 |
| ss2_TE_184 | LTR/Gypsy | 8826 | 1 | 1 | 7450 |
| ss2_TE_185 | LTR/Gypsy | 6698 | 5 | 5 | 8697 |
| ss2_TE_186 | LTR/Gypsy | 6709 | 1 | 1 | 6709 |
| ss2_TE_187 | LTR/Gypsy | 5661 | 4 | 2 | 5855 |
| ss2_TE_188 | LTR/Gypsy | 8335 | 1 | 1 | 8335 |
| ss2_TE_189 | LTR/Gypsy | 7997 | 3 | 2 | 8399 |
| ss2_TE_190 | LTR/Gypsy | 6684 | 2 | 2 | 6830 |
| ss2_TE_191 | LTR/Gypsy | 9953 | 4 | 2 | 10284 |
| ss2_TE_192 | LTR/Gypsy | 1447 | 9 | 5 | 6477 |
| ss2_TE_193 | LTR/Gypsy | 8335 | 4 | 2 | 8587 |
| ss2_TE_194 | LTR/Gypsy | 9944 | 9 | 3 | 10772 |
| ss2_TE_195 | LTR/Gypsy | 6688 | 1 | 1 | 6688 |
| ss2_TE_196 | LTR/Gypsy | 7972 | 10 | 6 | 7823 |
| ss2_TE_197 | LTR/Gypsy | 2777 | 2 | 1 | 2777 |
| ss2_TE_198 | LTR/Gypsy | 6691 | 3 | 3 | 7389 |
| ss2_TE_199 | LTR/Gypsy | 6710 | 2 | 2 | 12158 |
| ss2_TE_200 | LTR/Gypsy | 8281 | 3 | 1 | 7817 |
| ss2_TE_201 | LTR/Gypsy | 6683 | 3 | 3 | 12922 |
| ss2_TE_202 | LTR/Gypsy | 9263 | 12 | 9 | 14337 |
| ss2_TE_203 | LTR/Gypsy | 11495 | 6 | 2 | 11750 |
| ss2_TE_204 | LTR/Gypsy | 5950 | 18 | 13 | 13043 |
| ss2_TE_205 | LTR/Gypsy | 3060 | 63 | 55 | 40868 |
| ss2_TE_206 | LTR/Gypsy | 14935 | 28 | 3 | 17603 |
| ss2_TE_207 | LTR/Gypsy | 6711 | 2 | 2 | 13422 |
| ss2_TE_208 | LTR/Gypsy | 6326 | 1 | 1 | 5127 |
| ss2_TE_209 | LTR/Gypsy | 9795 | 1 | 1 | 9795 |
| ss2_TE_210 | LTR/Gypsy | 1982 | 1 | 1 | 1982 |
| ss2_TE_211 | LTR/Gypsy | 9931 | 1 | 1 | 9931 |
| ss2_TE_212 | LTR/Gypsy | 3288 | 2 | 1 | 1637 |
| ss2_TE_213 | LTR/Gypsy | 6697 | 2 | 2 | 7293 |
| ss2_TE_214 | LTR/Gypsy | 7759 | 4 | 3 | 8517 |
| ss2_TE_215 | LTR/Gypsy | 5531 | 3 | 2 | 5706 |
| ss2_TE_216 | LTR/Gypsy | 8004 | 3 | 1 | 8004 |
| ss2_TE_217 | LTR/Gypsy | 6683 | 1 | 1 | 6683 |
| ss2_TE_218 | LTR/Gypsy | 6706 | 5 | 4 | 16164 |
| ss2_TE_219 | LTR/Gypsy | 792 | 1 | 1 | 792 |
| ss2_TE_220 | LTR/Gypsy | 6710 | 3 | 3 | 11430 |
| ss2_TE_221 | LTR/Gypsy | 813 | 1 | 1 | 813 |
| ss2_TE_222 | LTR/Gypsy | 9956 | 8 | 6 | 28014 |
| ss2_TE_223 | LTR/Gypsy | 5662 | 7 | 2 | 5839 |
| ss2_TE_224 | LTR/Gypsy | 14753 | 15 | 8 | 26061 |
| ss2_TE_225 | LTR/Gypsy | 417 | 3 | 3 | 1249 |
| ss2_TE_226 | LTR/Gypsy | 5668 | 1 | 1 | 5667 |
| ss2_TE_227 | LTR/Gypsy | 5688 | 2 | 1 | 5688 |
| ss2_TE_228 | LTR/Gypsy | 6718 | 2 | 1 | 6718 |
| ss2_TE_229 | LTR/Gypsy | 8007 | 2 | 1 | 907 |
| ss2_TE_230 | LTR/Gypsy | 6696 | 2 | 2 | 7293 |
| ss2_TE_231 | LTR/Gypsy | 7998 | 16 | 11 | 31997 |
| ss2_TE_232 | LTR/Gypsy | 6651 | 4 | 3 | 13385 |
| ss2_TE_233 | LTR/Gypsy | 8186 | 18 | 9 | 14124 |
| ss2_TE_234 | LTR/Gypsy | 6709 | 4 | 4 | 10508 |
| ss2_TE_235 | LTR/Gypsy | 9136 | 2 | 1 | 8281 |
| ss2_TE_236 | LTR/Gypsy | 3302 | 4 | 1 | 2057 |
| ss2_TE_237 | LTR/Gypsy | 9131 | 3 | 1 | 9131 |
| ss2_TE_238 | LTR/Gypsy | 2001 | 4 | 4 | 7476 |
| ss2_TE_239 | LTR/Gypsy | 6711 | 1 | 1 | 6711 |
| ss2_TE_240 | LTR/Gypsy | 6720 | 5 | 3 | 7837 |
| ss2_TE_241 | LTR/Gypsy | 8010 | 3 | 1 | 8010 |
| ss2_TE_242 | LTR/Gypsy | 9136 | 7 | 1 | 9136 |
| ss2_TE_243 | LTR/Gypsy | 6326 | 10 | 8 | 9097 |
| ss2_TE_244 | LTR/Gypsy | 2778 | 1 | 1 | 2778 |
| ss2_TE_245 | LTR/Gypsy | 8516 | 60 | 17 | 19813 |
| ss2_TE_246 | LTR/Gypsy | 6684 | 1 | 1 | 6684 |
| ss2_TE_247 | LTR/Gypsy | 6689 | 5 | 3 | 7277 |
| ss2_TE_248 | LTR/Gypsy | 9944 | 6 | 6 | 18996 |
| ss2_TE_249 | LTR/Gypsy | 543 | 1 | 1 | 543 |
| ss2_TE_250 | LTR/Gypsy | 6699 | 2 | 2 | 10344 |
| ss2_TE_251 | LTR/Gypsy | 8028 | 16 | 5 | 17472 |
| ss2_TE_252 | LTR/Gypsy | 6697 | 1 | 1 | 6697 |
| ss2_TE_253 | LTR/Gypsy | 6326 | 2 | 1 | 4302 |
| ss2_TE_254 | LTR/Gypsy | 9670 | 5 | 3 | 29009 |
| ss2_TE_255 | LTR/Gypsy | 9964 | 5 | 3 | 16467 |
| ss2_TE_256 | LTR/Gypsy | 12925 | 26 | 10 | 22569 |
| ss2_TE_257 | LTR/Gypsy | 834 | 1 | 1 | 834 |
| ss2_TE_258 | LTR/Gypsy | 6682 | 1 | 1 | 6682 |
| ss2_TE_259 | LTR/Gypsy | 6400 | 1 | 1 | 6400 |
| ss2_TE_260 | LTR/Gypsy | 5563 | 3 | 1 | 5563 |
| ss2_TE_261 | LTR/Gypsy | 6683 | 2 | 2 | 13357 |
| ss2_TE_262 | LTR/Gypsy | 6684 | 1 | 1 | 6684 |
| ss2_TE_263 | LTR/Gypsy | 3597 | 20 | 7 | 15501 |

**Table S4.**
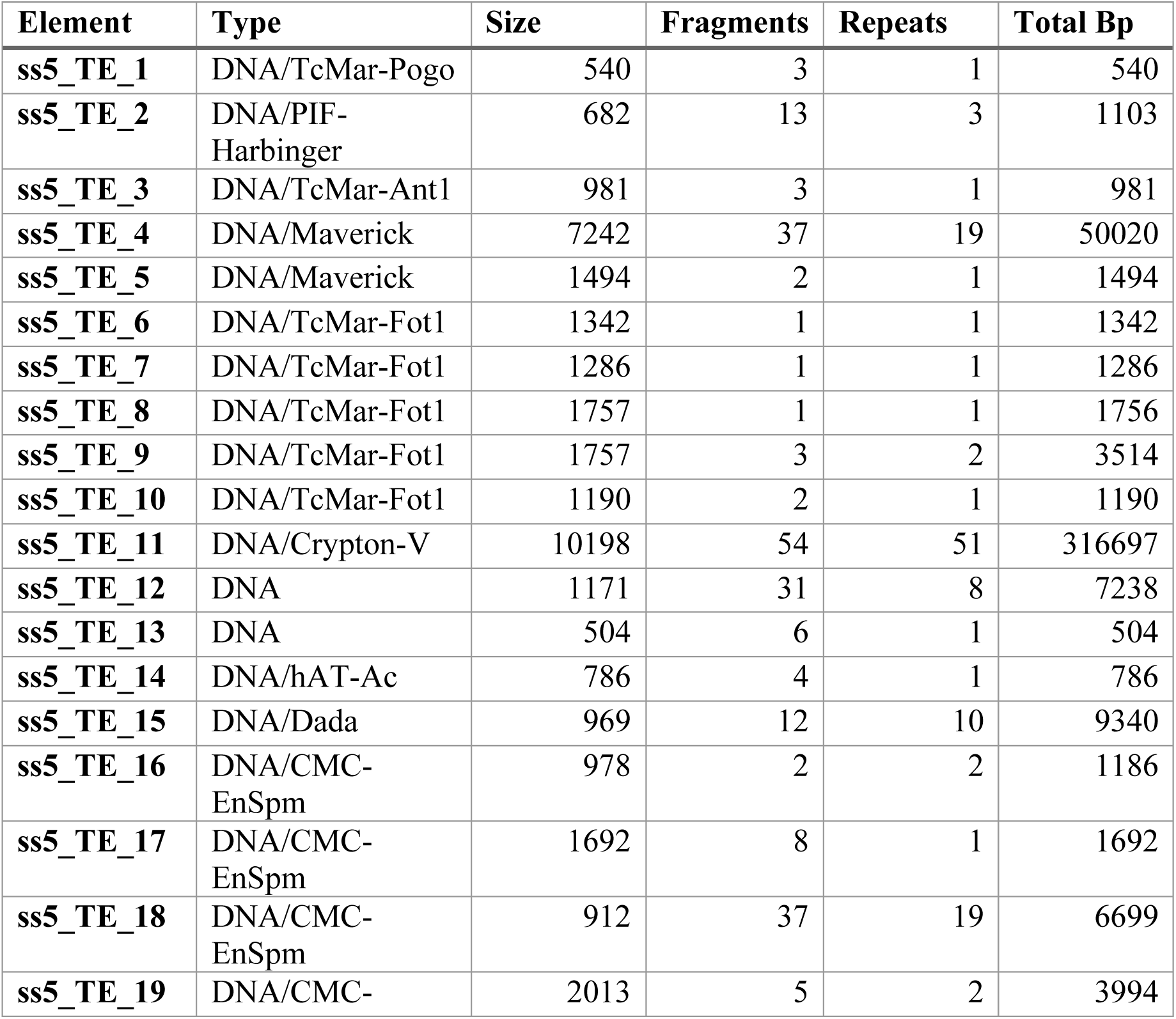
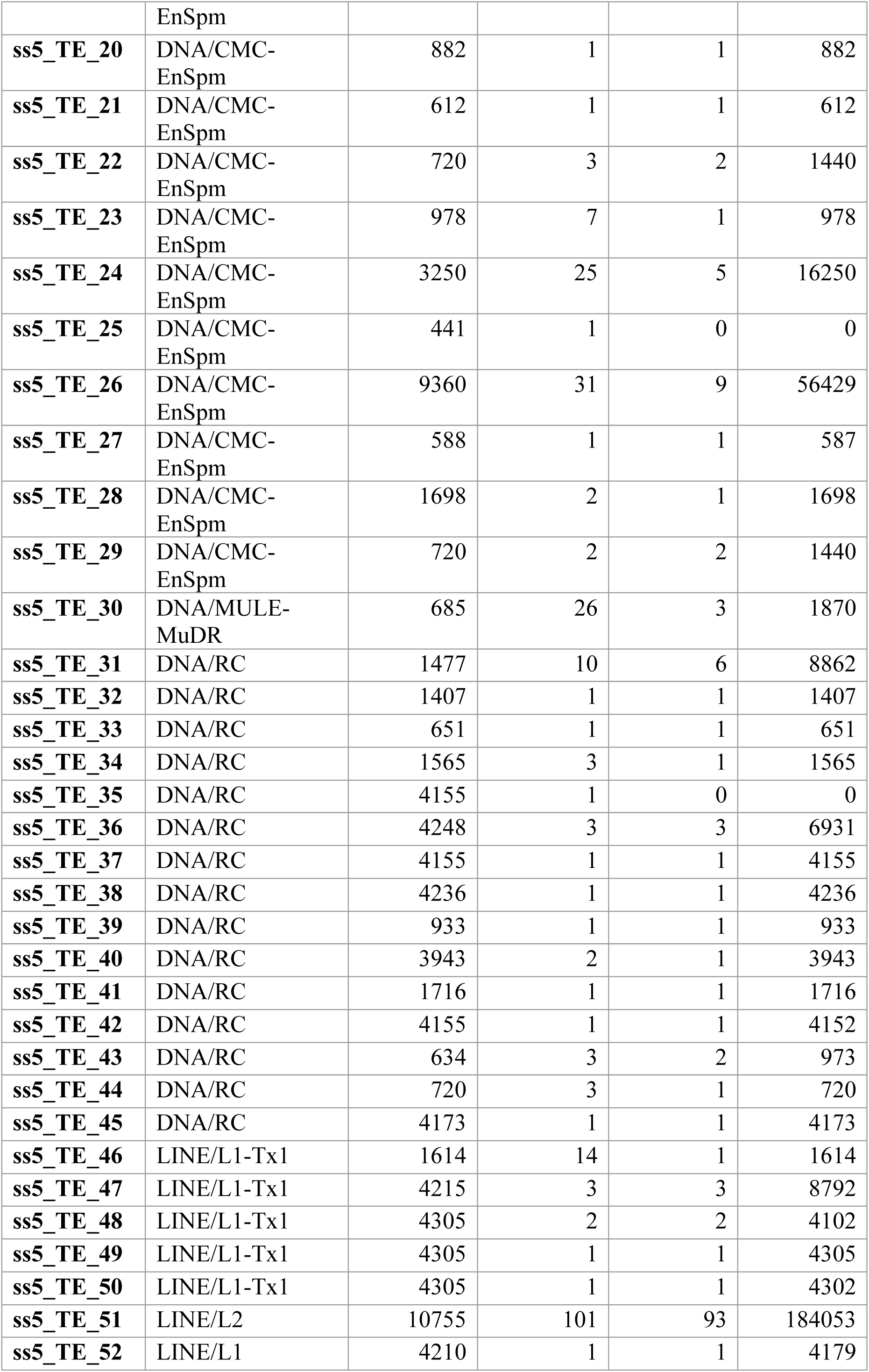
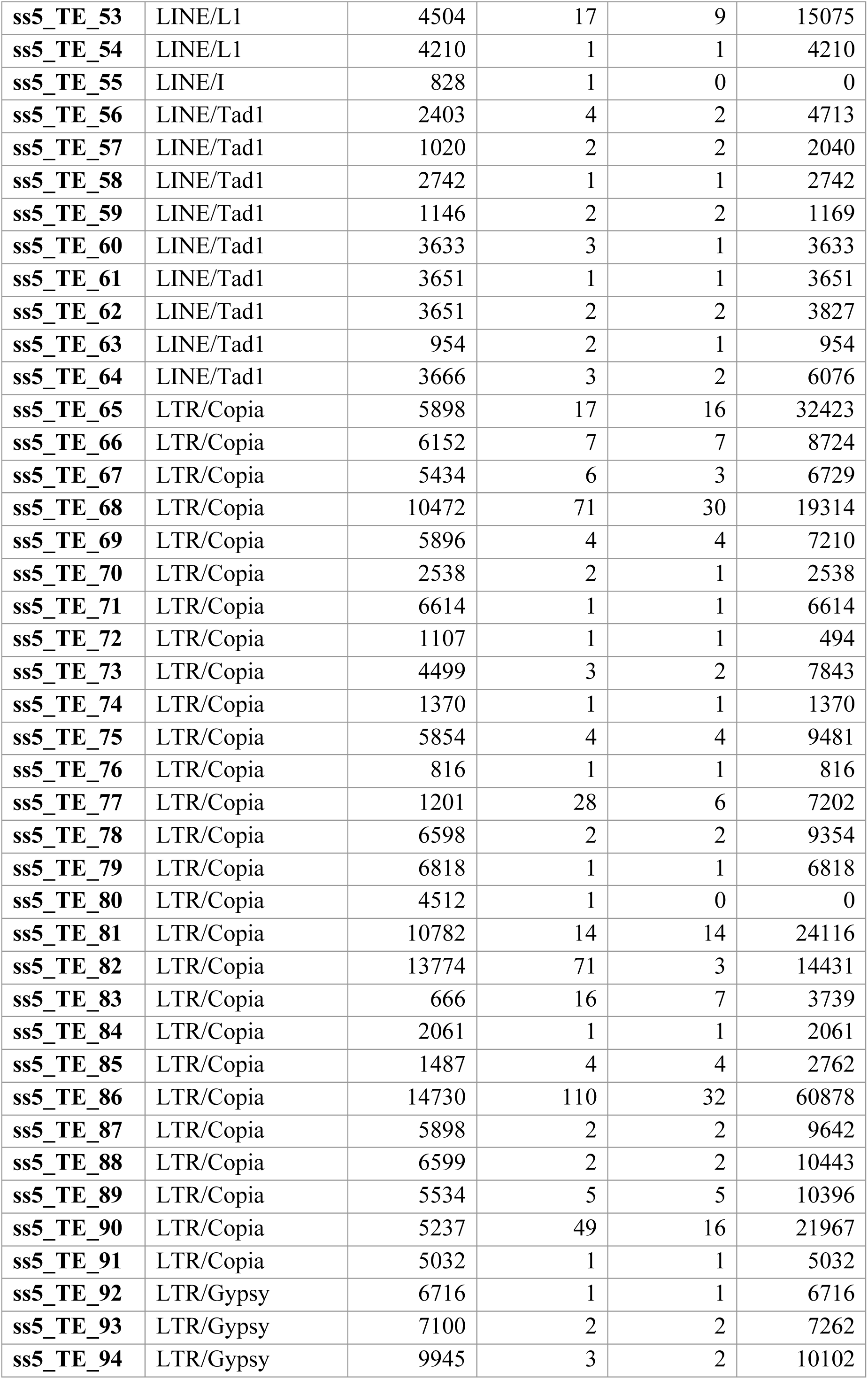
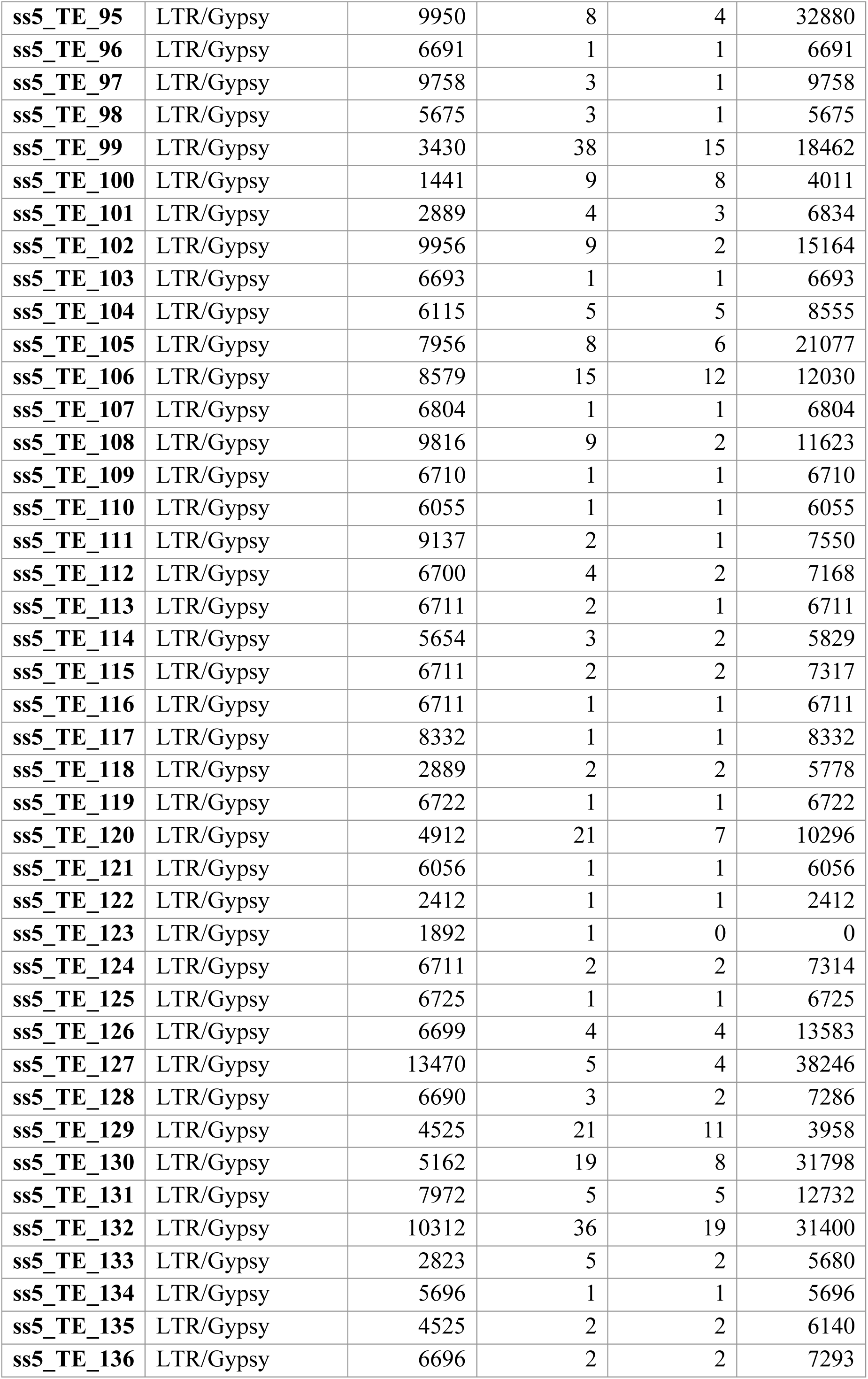
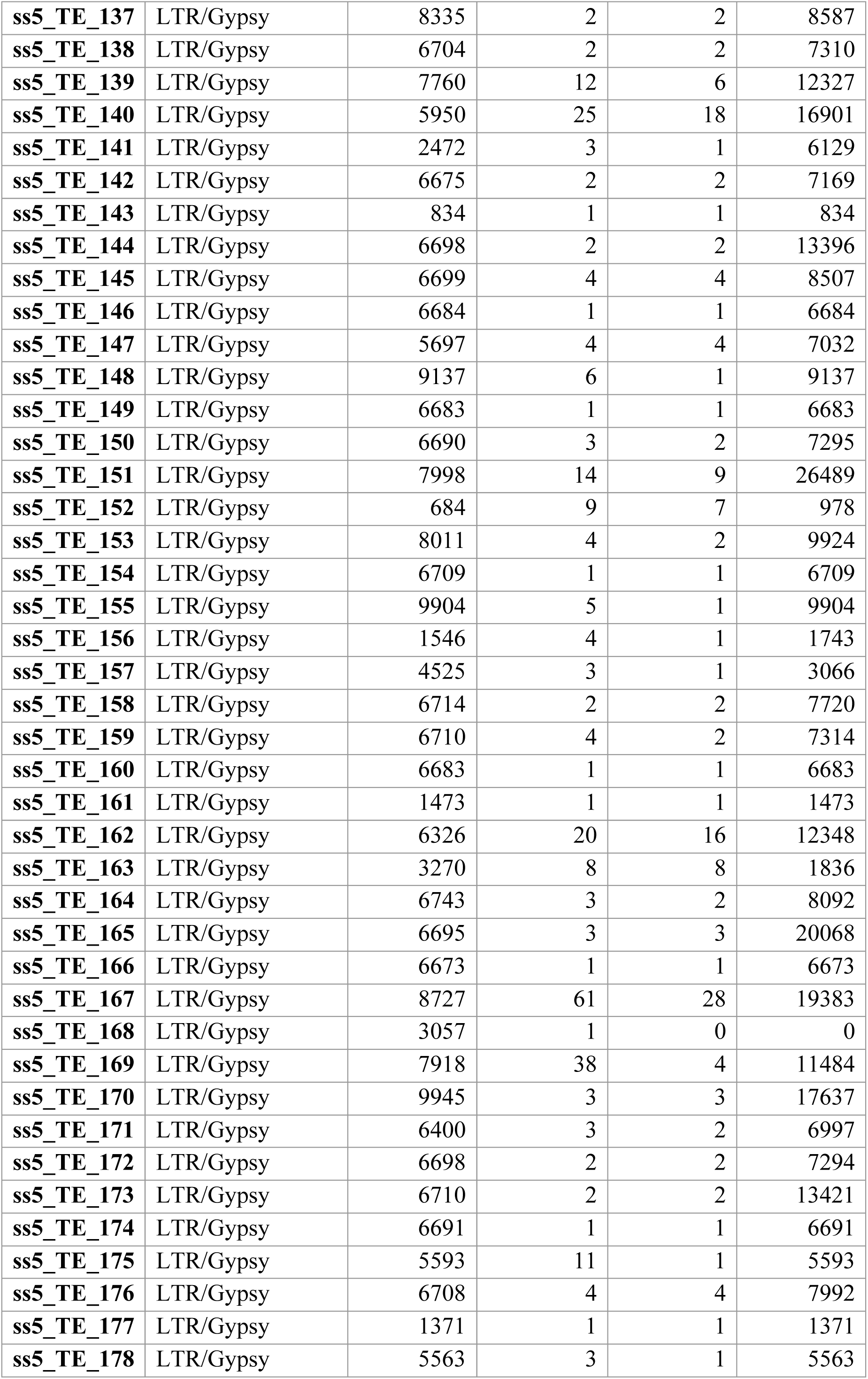
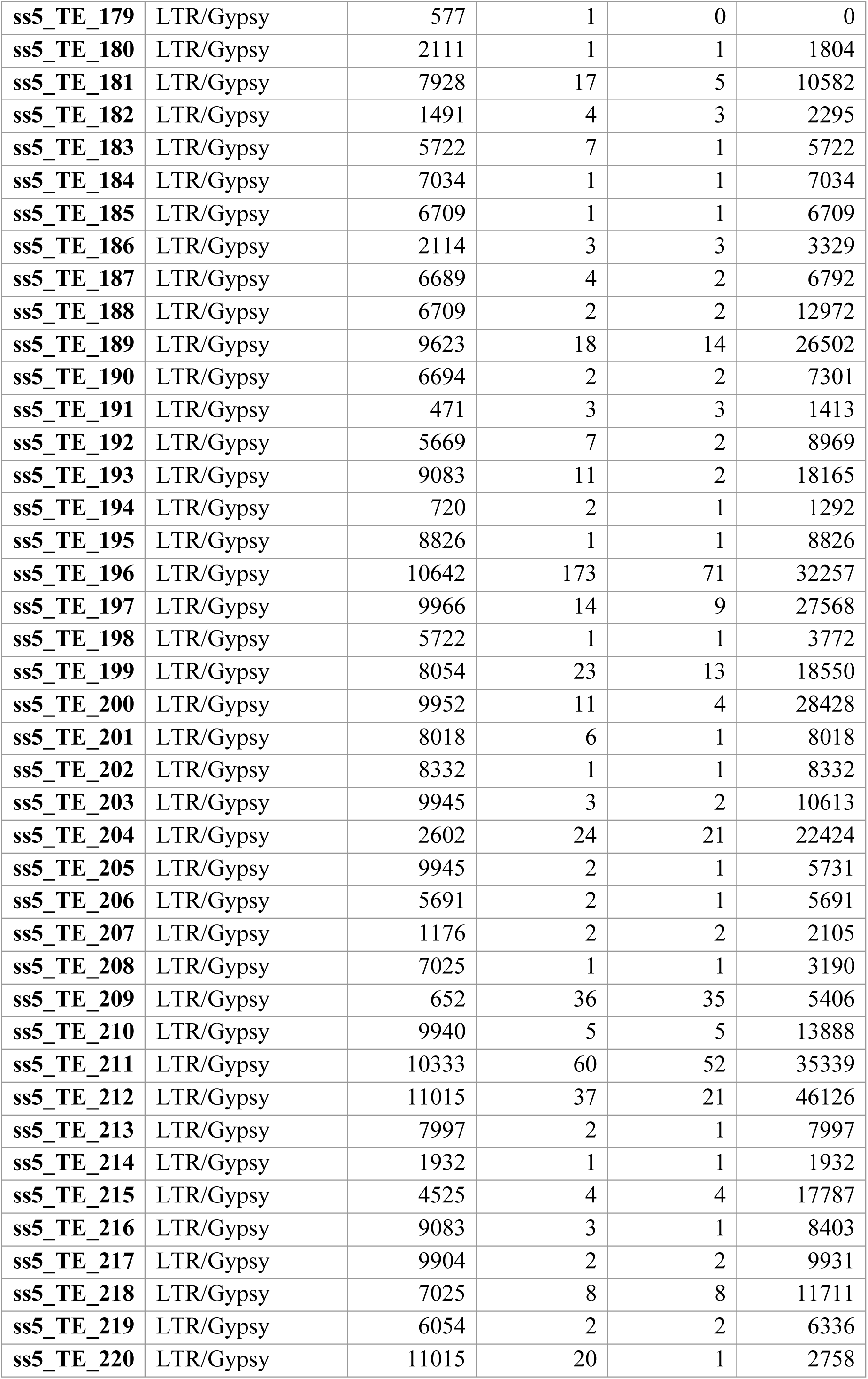
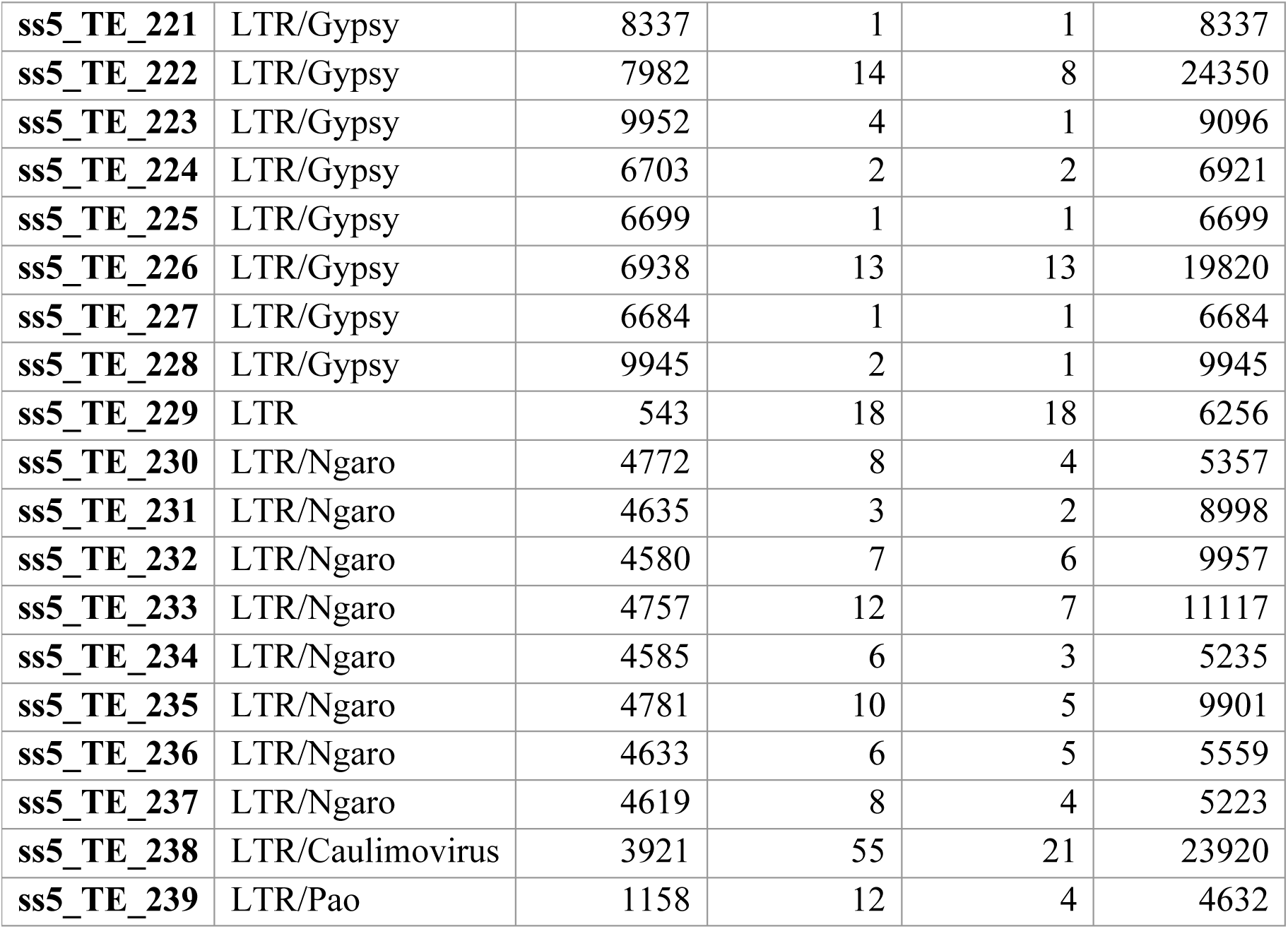
Detected TE elements in *P. pulmonarius* ss5.

**Table S5.**
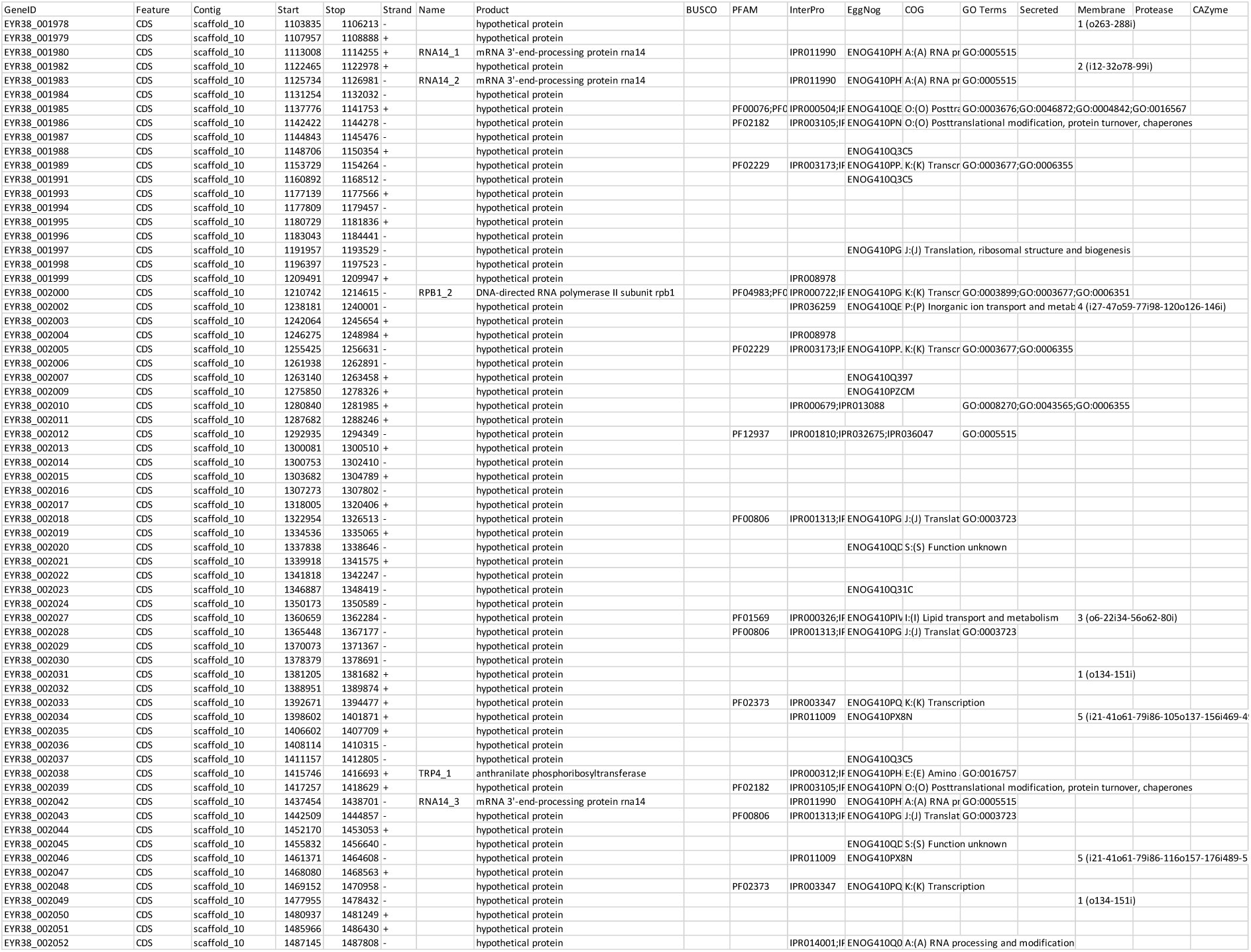
List of unique genes of ss2 (without homologs in ss5) clustered at the end of scaffold 10 together with their functional annotation.

**Table S6.**
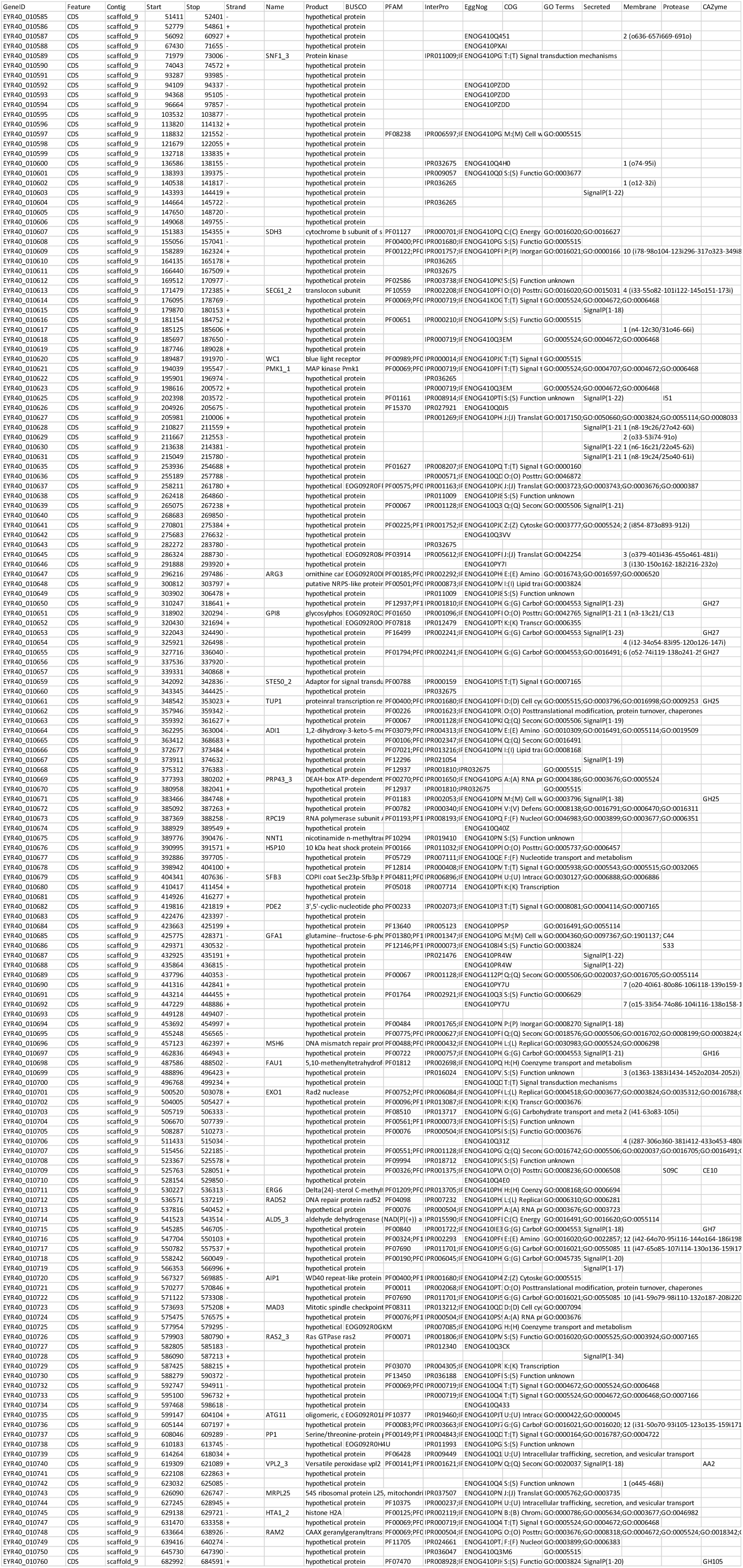
List of unique genes of ss5 (without homologs in ss2) clustered at the beginning of scaffold 9 together with their functional annotation.

